# Red blood cells protein profile is modified in breast cancer patients

**DOI:** 10.1101/2022.01.04.474889

**Authors:** Thais Pereira-Veiga, Susana Bravo, Antonio Gómez-Tato, Celso Yáñez-Gómez, Carmen Abuín, Vanesa Varela, Juan Cueva, Patricia Palacios, Ana B. Dávila-Ibáñez, Roberto Piñeiro, Ana Vilar, María del Pilar Chantada-Vázquez, Rafael López-López, Clotilde Costa

**Affiliations:** Roche-Chus Joint Unit, Translational Medical Oncology Group, Oncomet, Health Research Institute of Santiago de Compostela (IDIS), Travesía da Choupana s/n, 15706 Santiago de Compostela, Spain; Department of Tumor Biology, Center of Experimental Medicine, University Medical Center Hamburg-Eppendorf, 20246, Hamburg, Germany; Proteomic Unit, Instituto de Investigaciones Sanitarias-IDIS, Complejo Hospitalario Universitario de Santiago de Compostela (CHUS), 15706 Santiago de Compostela, Spain; CITMAga, University of Santiago de Compostela (Campus Vida), 15782 Santiago de Compostela, Spain; Department of Oncology, University Hospital of Santiago de Compostela (SERGAS), 15706 Santiago de Compostela, Spain; CIBERONC, Centro de Investigación Biomédica en Red Cáncer, 28029 Madrid, Spain; Department of Gynecology, University Hospital of Santiago de Compostela (SERGAS), Spain

**Keywords:** Red Blood Cells (RBCs), breast cancer, metastasis, LAMP2

## Abstract

Metastasis is the primary cause of death for most breast cancer patients who succumb to the disease. During the haematogenous dissemination, circulating tumor cells interact with different blood components. Thus, there are micro-environmental and systemic processes contributing to cancer regulation. We have published that Red Blood Cells (RBCs) that accompany circulating tumor cells have prognostic value in metastatic breast cancer patients. Although the principal known role of RBCs is gas transport, it has been recently assigned additional functions as regulatory cells on circulation. Hence, to explore their potential contribution to tumor progression, we characterized the proteomic composition of RBCs from 53 breast cancer patients, compared with 33 healthy donors. RBCs from breast cancer patients showed a different proteomic profile compared to healthy donors. The differential proteins were mainly related to extracellular components, proteasome, and metabolism. Besides, LAMP2 emerge as a new RBCs marker with diagnostic and prognostic potential for metastatic patients. Seemingly, RBCs are acquiring modifications in their proteomic composition that probably represents the systemic cancer disease, conditioned by the tumor microenvironment.

## Introduction

Breast carcinoma (BC), is the leading cancer-related cause of death in women and has a higher incidence rate than any other type of tumor. The metastatic disease accounts for the overwhelming majority of cancer-related deaths, thus, the lack of reliable biomarkers for early metastasis diagnoses remains as one of the main clinical challenges. Tumor metastasis involves a multistep process where cancer cells escape from their primary site, circulate in the bloodstream as circulating tumor cells (CTCs), and then extravasate through the vascular walls into the parenchyma of distant tissues, where they finally adapt and outgrowth(1). Once CTCs reach and settle in a distant organ they are called disseminated tumor cells (DTCs) and together with CTCs are recognized as the seeds of metastasis(2). Interactions between CTCs and normal blood components such as platelets, neutrophils, monocytes, and endothelial cells are crucial for their survival in the bloodstream and can facilitate the extravasation at distant sites(3). Surprisingly, little is known about the role of the most abundant component of the blood, the red blood cells (RBCs), in the metastatic process.

RBCs account for nearly > 70% of the total cell count in the average adult(4). Erythropoiesis begins with the differentiation of multipotent hematopoietic stem cells in the bone marrow, which then give rise to erythroid-committed precursors. In the last stages of the process, the nucleus and other organelles are extruded, and these enucleated reticulocytes are released into the bloodstream to complete their maturation process in a tightly regulated process. Related to erythropoiesis, abnormal Red blood cell Distribution Width (RDW), has been associated with poor prognosis in cancer(5) and advanced disease(6,7). Our group has recently reported that the presence of escort RBCs in the enriched CTCs fraction was linked with a worse outcome on metastatic BC patients(8). This observation suggested alterations in the RBCs of patients with metastatic BC.

This study aimed to provide for the first time a large-scale proteomic analysis of RBCs from healthy donors (HD) and BC patients. We extensively analyzed HD and treatment-naïve metastatic and non-metastatic BC patients using two proteomic analyses (shotgun, and SWATH-MS), to identify potential differences in proteomic profiles in BC patients’.

## Results

### Differential Red Blood Cells proteomic profiles between Breast Cancer patients and Healthy Donors

The proteomic analysis by liquid chromatography-tandem mass spectrometry (LC-MS/MS) in data-dependent acquisition (DDA) mode-Shotgun of RBCs of samples from BC patients (non-metastatic: M0, n= 17; metastatic: M1, n=19 and HD, n=21) mainly reported specific proteins of RBCs such as haemoglobins (Hbs) or spectrins (Figure 1A, Table S2). However, the RBCs proteomic profile of BC patients resulted to be different from HD.

**Figure 1.**
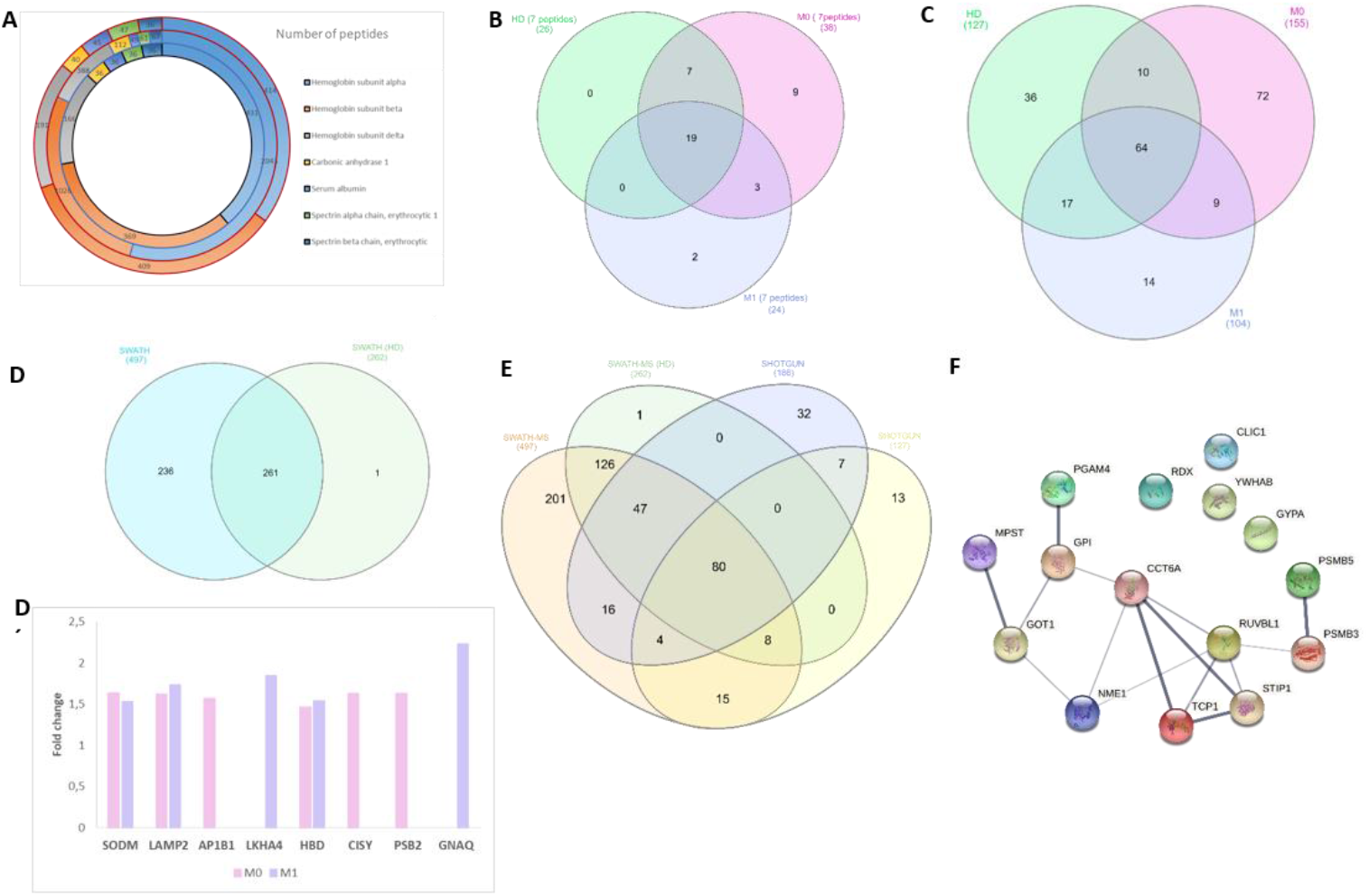
Massive proteomic analysis of RBCs from BC patients and HD. A) Representation of the number of major peptides detected by shotgun in healthy donors (HD) (the inner black circle), M0 patients (the intermediate blue circle), and M1 patients (the external red circle). B-C) Venn diagram showing the common proteins among patients (M0 and M1) and HD RBCs samples identified using shotgun technology for ≥ 7 peptides (B) and ≥ 1 peptide (C). D) Venn diagram showing the common proteins among BC patients and HD RBCs samples identified using SWATH-MS technology. D’) Relative fold change of M0 and M1 compared to HD of the indicated proteins. E) Venn diagram showing the common proteins among both approaches. F) String network analysis showing interactions between 16 common proteins identified by both approaches exclusively of BC patients.

We identified 14 proteins as unique to the patient samples when considering ≥ 7 peptides per protein and FDR < 1% (Figure 1B, Table S2). These proteins were associated with the metabolism of amino acids, glycolysis, and gluconeogenesis by Kegg pathways (PGK1, BPGM, ALDOA, and TPI1 proteins) (PPI enrichment p-value: 4.4e-07). In addition, the potentially related diseases to specific patients’ proteins were anemias or haematopoietic system diseases (proteins HBZ, EPB41, HP, PGK1, HBE1, BPGM, and ALDOA). Embryonic Hbs isoforms such as epsilon (HBE1) and zeta (HBZ) were identified exclusively in M1 and M0 patients, respectively.

Next, we checked all the detected proteins (≥ 1 peptide and FDR < 1%) comparing HD and BC patients (Figure 1C, Table S3-5). M0 BC patients showed 72 differential proteins compared to HD, among the 260 proteins that comprise the library. In M1 patients, 14 proteins were identified exclusively to the advanced stage. Nine proteins were shared between M0 and M1 BC patients. Considering the 95 proteins unique to the patients’ cohort, GO analysis pointed to proteasome and chaperone complex (Kegg pathways)(PPI enrichment p-value < 1.0e-16). Regarding the biological processes, regulation of amino acid metabolism or extracellular exosomes were highlighted. On these samples were also found proteins that are usually localized in Cajal bodies. No specific proteins of platelets, neutrophils, or lymphocytes were found on this analysis, proving the absence of other blood cells in the RBCs fraction of the analyzed samples. Besides, no erythroid precursor markers were observed.

Seeing that the RBCs of patients showed a different proteomic profile, we took advantage of the Sequential Window Acquisition of All Theoretical Mass Spectra (SWATH-MS) strategy that enables quantitative analysis of proteins with high precision and consistency. The data output from HD RBCs samples (n=33) identified a total of 262 proteins (Table S6). The GSEA confirmed that the main enriched pathway was *erythrocytes take up carbon dioxide and release oxygen*. In agreement, one of the preferred tissues was RBCs.

Similar to shotgun data, proteomic profile from BC patients’ RBCs (M0, n= 26; M1, n=27) was different to HD (n=33) by SWATH-MS (t-test, p < 0.05) (Figure 1D). The generated library including patients and HD comprised 497 RBCs proteins. Proteins found only in BC patients were related to integrin and inflammatory signalling pathways or extracellular matrix interaction and platelet degranulation. Next, we performed the quantitative analysis comparing HD to both M0 and M1 BC patients. Compared to HD, in M1 samples, 22 proteins were upregulated 15 proteins were down-regulated while in M0 samples, 10 proteins were upregulated and 17 were down-regulated (Table S7). Figure 1D’ depicted those proteins that showed a greater fold change of expression in BC patients compared to controls, as GNAQ, LKHA4, or LAMP2. The GSEA of the proteins differentially expressed between patients and HD emphasized the pentose phosphatase, biosynthesis of amino acids and, secretory granule lumen pathway (PPI enrichment p-value: < 1.0e-16). In addition, proteins linked to congenital haemolytic anemia were also identified in these samples. Among the proteins differentially expressed in patients, no proteins specific to RBC precursors or other white blood cells were identified. BC patient’s samples also showed differential protein expression based on the stage (M0 or M1). Thus, 10 proteins were up-regulated in M1 compared to M0, while 7 proteins were down-regulated (Table S7). Of those, some proteins were also altered versus HD (Table S7). These proteins were also found in congenital haemolytic anemia (PPI enrichment p-value: 0.000359).

#### Comparative analysis between shotgun and SWATH-MS proteomics data identified common proteins

The proteins identified by *shotgun* and SWATH-MS in these samples showed a high overlap, founding 79 % of identified proteins by *shotgun* also in SWATH-MS data (Figure 1E). The GO analysis indicated similar pathways or biological processes, reinforcing the similarity in both approaches’ data. Of the proteins exclusive of BC patients, 16 matched in both approaches. These proteins are related to the chaperone complex and carbon metabolism (PPI enrichment p-value: 2.36e-06) (Figure 1F).

### Identification of novel proteins in red blood cells

The lists of identified proteins from HD donors in this study by shotgun and/or SWATH-MS approaches were compared with the Uniprot database for RBCs and the database Repository of Enhanced Structures of Proteins Involved in the Red blood cell Environment (RESPIRE). This latter database aims to provide a comprehensive reference of protein-based information on the proteins available in the RBCs. In addition, our data were compared with repositories from other publications(9,10). In total, 50 new proteins not previously described in RBCs databases were found in this study (Figure 2A, A’, Table S8). These proteins were mainly linked to neutrophil degranulation and phagosome by GSEA (PPI enrichment p-value: < 1.0e-16) (Figure 2B).

**Figure 2.**
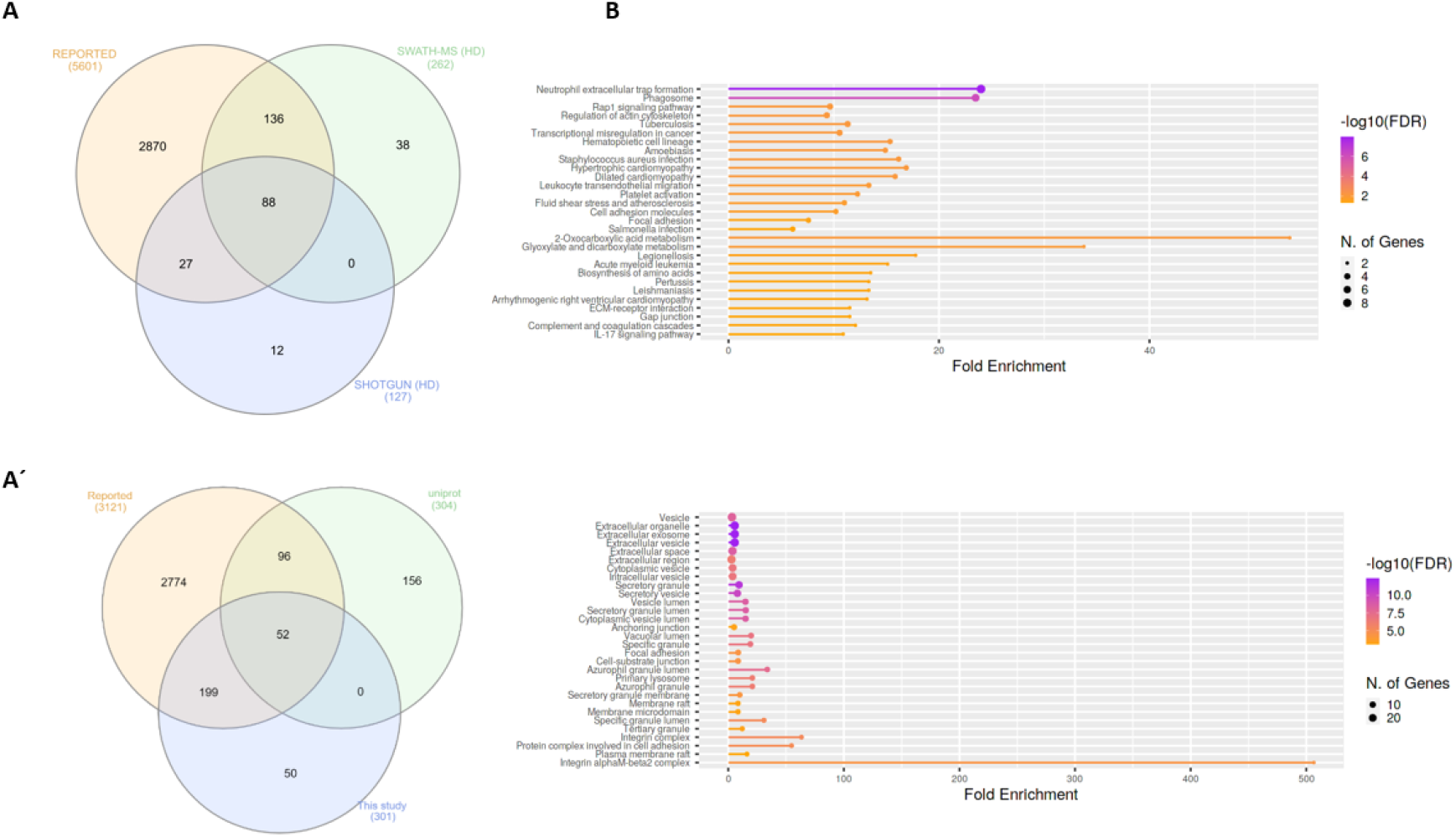
Identification of novel RBCs proteins in healthy donor samples. A, A’) Venn diagram showing the common proteins between this study (shotgun or SWATH-MS) and the reported proteins including the RESPIRE project (https://www.dsimb.inserm.fr/respire/) and two related publications (Bryk et al and Alessandro et al) (9,10)(A) or Uniprot database (A’). B) GO of the 50 novel proteins identified regarding the KEGG pathway and Cellular components (ShinyGO v0.741).

### Metastatic breast cancer patients showed altered blood clinical values

Blood cells indexes are included in all standard clinical tests. To check whether blood clinical parameters can be affected by the presence of BC or the tumor stage, these values were compared between the HD and patients included in this study. Haematocrit and Haemoglobin (Hb) concentrations were significantly lower in the M1 cohort (p< 0.0001, Kruskal-Wallis test) (Figure 3 A, B). A slight increase of monocytes counts was observed on the M1 patients compared with HD (p=0.07, Mann Whitney test) (Figure 3C) while no differences were found for the count of lymphocytes, eosinophils, basophils, neutrophils, or platelets between the three groups. Interestingly, the levels of blood parameters of M0 patients were closer to HD than to M1 BC patients.

**Figure 3.**
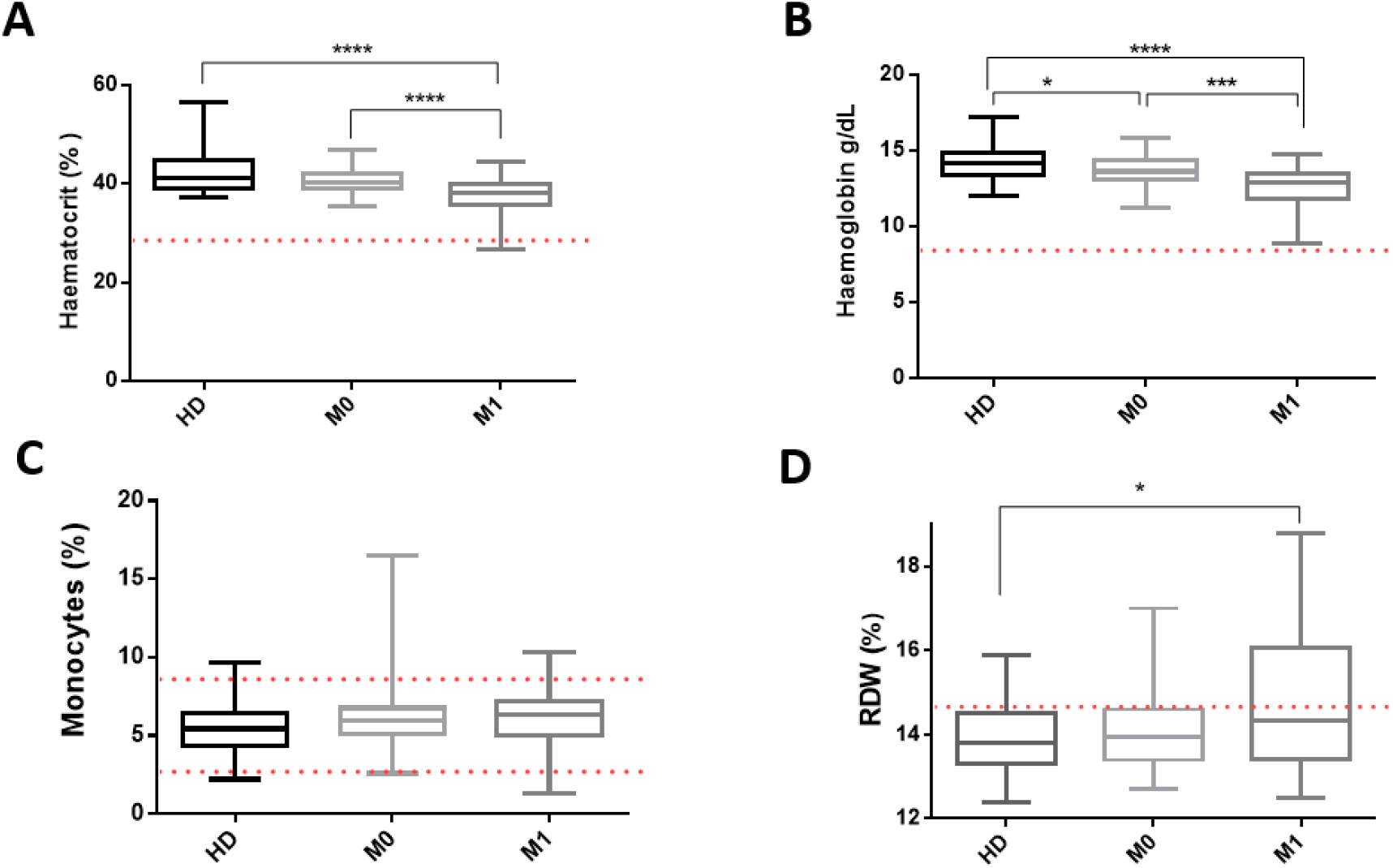
Altered blood test parameters in non-metastatic (M0) (n=44) and metastatic (M1) (n=34) Breast Cancer (BC) patients compared with healthy donors (HD) (n=30). **A)** The hematocrit represents the percentage of RBCs. The lower limit is 36.9 %. M1 BC patients had a lower hematocrit compared to HD or M0 patients. **B)** Concentration of Hb in the blood. The lower limit is 12.2 g/dL. M1 BC patients had a lower concentration compared to HD or M0 patients. Besides, M0 patients showed a lower level compared to HD. **C)** The standard percentage of monocytes oscillates between 2.7-8.6. M1 BC patients have a slight increase in monocytes compared with HD, close to statistical significance. Only 6 patients had altered parameters from standard ones. **D)** Red Distribution Width (RDW) represents the percentage of Red blood cells with abnormal size. The standard range is 11.5 to 14.5. M1 BC patients presented a high value compared with HD. The red dotted line represents the limit of the standard value for each parameter. P-value < 0.05 (*); < 0.01 (**) and < 0.001 (***).

The RDW parameter, which measures the variation in the volume and size of RBCs, was altered in the advanced disease. Thus, M1 patients showed higher values (mean 14.96 ± 1.92) compared with the M0 (mean 14.07 ± 0.93) or the HD group (mean 13.88 ± 0.77) (Figure 3D),(p=0.01, Mann Whitney test). Contingency analysis considering elevated (> 14.5) or normal RDW (≤ 14.5) indicated that altered RDW was associated with M1 patients (p=0.002, Fisher exact test).

### LAMP2 as a prognostic and diagnostic marker for metastatic BC

To check if the protein levels determined by SWATH-MS were linked with the patient’s outcome, those proteins with a fold change higher than 1.5 in M1 patients versus HD samples were selected (Table S7). To decipher the predictive potential of GNAQ, HBD, LKHA4, and LAMP2, a survival analysis was performed. The high or low protein expression levels were determined using the percentile 70. Protein levels of GNAQ, HBD, or LKHA4 did not show prognostic value in this cohort of study (log-rank test p > 0.05, n=27). However, high LAMP2 expression levels can predict the worst outcome in M1 BC patients (PFS: p= 0.0005, 3.5 vs 23.73 months, log-rank test; OS: p=0.05, log-rank test) (Figure 4 A, B). Besides, low levels of LAMP2 were found in patients without disease progression (p=0.02, Chi-square test), or who were still alive during follow-up (p=0.03, Fisher’s exact test). Low LAMP2 levels were also associated with having normal RDW value (p=0.02, n=65: 18 CT, 22 M0, and 25 M1 samples). Altogether, the enhancement in LAMP2 levels was related to advanced disease. Interestingly, the purine nucleoside phosphorylase (PNP), whose specific enzymatic activity is very high in RBCs, showed lower levels in patients with bone metastases (p = 0.02, Fisher’s exact test).

**Figure 4.**
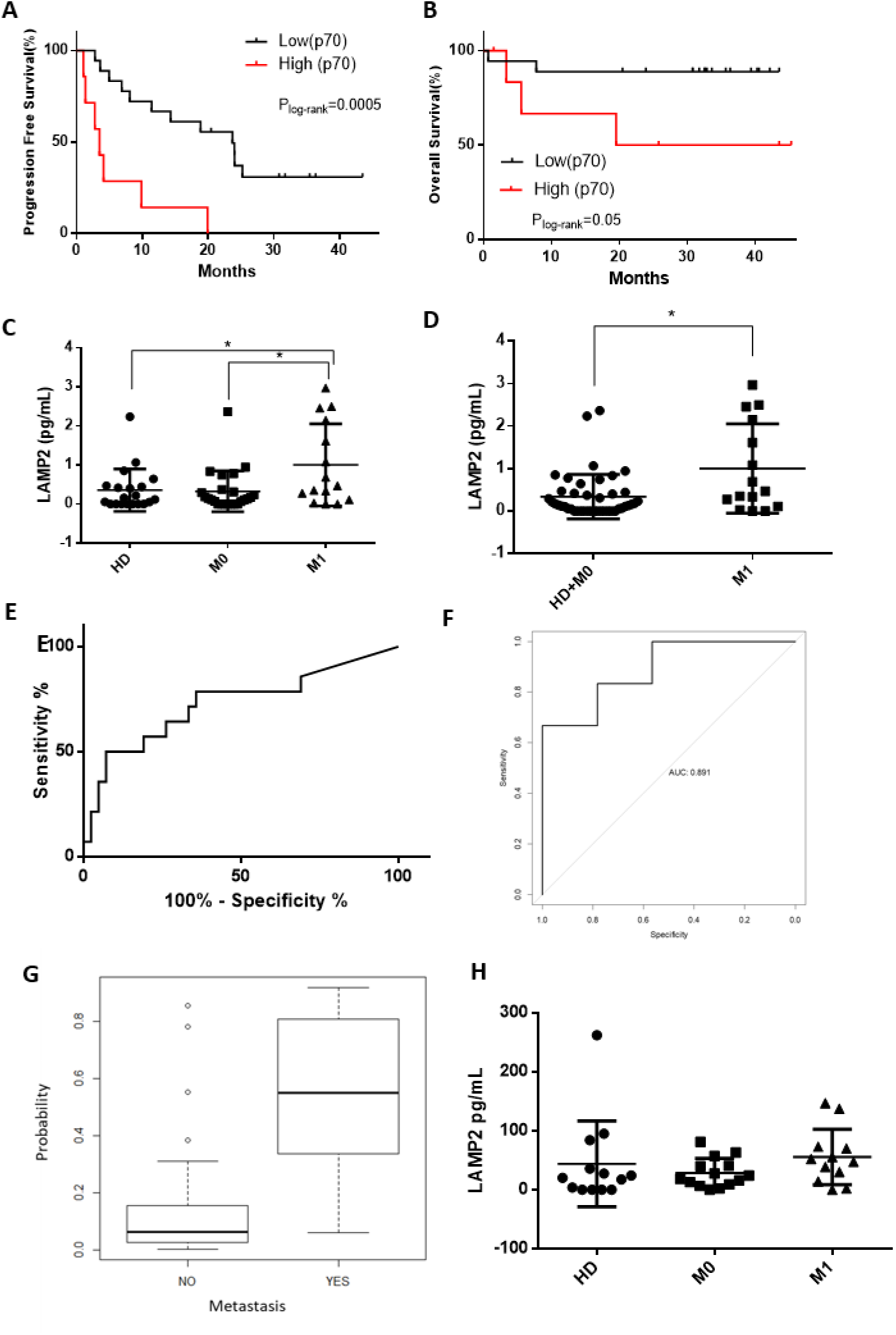
LAMP2 as a prognostic and diagnostic marker for metastatic BC. A, B) Kaplan-Meier plot for PFS (A) and OS (B). Percentile 70 defines high or low levels of LAMP2. C, D) LAMP2 concentration levels from RBCs by ELISA assay in healthy donors (HD, n=20), non-metastatic (M0, n= 24) and metastatic (M1, n=15) (C), or non-metastatic (M0 and HD, n=44) and metastatic (M1, n=15) (D). Non-metastatic patients were recruited pre-surgery. HD samples were obtained from women with paired age to the patients. E) ROC Curve for LAMP2 concentration levels. (AUC= 0.71). F) ROC Curve for the predictive model using LAMP2+ Haematocrit and RDW (AUC= 0.89). G) Probability of being metastatic given by the model. H) LAMP2 concentration levels from plasma by ELISA assay in HD, M0, and M1 (n=13, each group).

LAMP2 is a lysosome-related membrane glycoprotein that has been associated with BC tumor cells. However, its function in RBCs is unknown. An increase in LAMP2 levels in RBCs could indicate a systemic alteration in advanced disease and therefore may have diagnostic value. For that, LAMP2 expression was further validated by ELISA in RBCs lysates. As shown in Figure 4C, LAMP2 expression level on M1 samples was significantly higher than HD or M0 samples (p=0.04 and 0.02 respectively, Mann Whitney test), whereas no differences were observed between HD and the M0 cohort. Additionally, M1 samples showed higher LAMP2 expression levels when compared with all the other samples together (HD or M0 samples)(p=0.01, Wilcoxon test)(Figure 4D). Next, to evaluate LAMP2 levels as a diagnostic test for metastatic BC, Receiver-Operating Characteristic Analysis was performed including healthy donors (n=20), non-metastatic (M0, n= 24), and metastatic patients (M1, n=15). The area under the curve (AUC) value was 0.71, with 76 % sensitivity and 62 % specificity, thus LAMP2 expression levels of the RBCs were able to discriminate metastatic BC patients (Figure 4E). Besides, to obtain a more robust predictive model, those blood test parameters that were altered in metastatic BC patients, as RDW and haematocrit, were included. This logistic regression model increased the discriminatory potential of metastatic BC patients (sensitivity = 92.3 %, specificity = 80.5 %, AUC = 0.89) (Figure 4F), with a total success rate of 83.33 % (Figure 4G).

Since LAMP2 is ubiquitously expressed in white blood cells, plasma samples were also tested by ELISA assay. No differences were found between BC patients and HD in LAMP2 protein levels in plasma, proving the specific RBCs origin of the increased LAMP2 expression (Figure 4H).

## Discussion

RBCs are traditionally considered exclusively as gas transporters. However, the constant renewal of circulating RBCs consumes large amounts of energy daily, suggesting a central role for RBCs in human physiology and homeostasis(11). RBC’s average life span is 120 days, which may reflect the systemic imbalance in the body. Consequently, they act as markers of specific diseases and their evolution(12). Altered RBCs have been described in several diseases such as diabetes and Alzheimer’s, multiple sclerosis(13), and rheumatoid arthritis(14). Furthermore, the crosstalk between RBCs and immune cells has been described to cause progression of atherosclerotic disease(15). More recently, RBCs have been described to bind cell-free DNA, which leads to phagocytosis of RBCs and innate immune activation in pathological settings(16), uncovering a previously unappreciated role of RBCs as critical players in inflammation. Furthermore, it has been reported that SARS-CoV-2 infection has a significant impact on RBC’s structural membrane homeostasis at the protein and lipid levels(17). In cancer patients, small-scale studies have been published, pointing to alterations in RBCs(18–20) or interaction with tumor cells(21). In addition, a study in children with neuroblastoma, described altered erythropoiesis, suggesting that primary tumor cells produce effects at distance and that the observed impairment could be mediated by extracellular vesicles released by the tumor cells(22). However, no reported studies are published using massive proteomics analysis. Hence, the major finding of this work is that RBCs from BC have a differential proteome profile compared with HD and it also varies among patients with different tumor stages.

GSEA analysis identified proteins exclusive of the BC samples related mainly to proteasome, exocytosis, and amino acid metabolism, along with neutrophil degranulation and exocytosis, phagosome, and binding.

Proteasome deregulation has been linked with aging and disease. In the context of cancer, altered proteostasis promotes cell survival and tumor growth, leading to the deregulation of various cellular pathways(23). Proteasome activity is well described during erythropoiesis(24) but also identified in mature RBCs(25). Dekel *et al*. demonstrated that *Plasmodium falciparum* parasites, cultured in fresh blood from a human donor, secrete extracellular vesicles that contain functional 20S proteasome complexes and that these can reshape the cytoskeleton of naïve RBCs(26). The amino acid metabolism, which is altered in many types of cancer has been linked as a hallmark of malignancy(27). Thus, amino acids increase the metabolic rate of tumor cells and promote survival and proliferation(28). It has been reported that the levels of amino acids in RBCs are higher than the corresponding serum levels(29). Furthermore, the existence of an amino acid exchange between RBCs and different tissues or their ability to quickly absorb and release amino acids has been proved(30), supporting the role of RBCs as amino acid transporters between organs. Thus, the presence of high concentrations of amino acids in RBCs without being required by internal metabolic processes could be explained if they serve as a metabolic supply to tumor cells.

In this study, it was shown the presence of higher embryonic/fetal Haemoglobins (Hbs) such as epsilon (HBE1), theta (HBAT) or zeta, (HBZ) in patients with BC. These Hbs correspond to < 1% of the total in adult mammals. Different types of Hbs(including HBE1 and HBZ) have been detected in glioblastoma cell lines(31) while HBE1 has been associated with radioresistance in colorectal cancer cells(32). Furthermore, tumor cells can directly generate erythroid cells, composed predominantly of embryonic Hbs, to obtain oxygen in response to hypoxia(33). Indeed, embryonic or fetal Hbs have a higher oxygen affinity(34), which could give them an advantage in oxygen transfer in conditions of hypoxia or anemia, conditions frequently seen in cancer patients(35,36). In CTCs, Haemoglobin beta (HBB) has been related to cell survival(37). Through proteomics approaches, higher levels of the different HB chains (including HBB), have been observed in BC patients. However, this is contrary to what we observed in the blood test, in which haemoglobin is lower in patients, especially in metastatic BC patients. The blood test determines total HB levels and does not provide information on whether there is an imbalance of the various HB chains that make up HBA, HBA2, or HBF. Besides, metastatic BC patients showed lower haematocrit compared with HD or non-metastatic BC, confirming previously reported works(38). The alteration of blood parameters can have a multifactorial origin, including treatment. M0 samples were obtained before surgery or neoadjuvant therapy while M1 samples were collected before therapy initiation. Thus, the potential influence of treatment is negligible. Hence, this blood alteration could boost the stress in erythropoiesis on the bone marrow and derive it to extramedullary erythropoiesis (EE) in other organs. EE is a compensatory mechanism that occurs in adults as a consequence of inadequate spinal function caused by haematological disorders, chronic infections, and cancer, which causes erythropoiesis to occur mainly on the spleen and the liver(39). Although the presence of erythrocytes containing HbF could be associated with EE, it has been described that HbF is not restricted to fetal erythropoietic organs, and it does not necessarily correlate with the process of EE(40). In addition, we found high values of RDW in the metastatic BC patients, in accordance with a reported meta-analysis that has shown that RDW may be a potential prognostic marker in cancer patients(41). Likewise, it has been suggested that an increment on immature RBCs in the circulation could be the underlying reason behind the rise in the RDW value in cancer patients.

However, there is some controversy, since some authors support the relationship between RDW and cancer is a reflection of the effect that inflammation and oxidative stress cause on RBCs and that are also cancer risk factors. If the erythropoiesis was severely affected, it would be expected to find an increase in erythroid precursors in peripheral blood. However, we did not find classical progenitor protein markers differentially expressed in BC patients. In this regard, HBD has been involved in the regulation of fetus-adult Hb switch(42) and was detected in hematopoietic stem cells and hepatocarcinoma cells(43,44). HBD was identified by both proteomic approaches, with an increase in protein levels both in non-metastatic and metastatic BC patients. In agreement with impaired erythropoiesis due to the presence of tumor cells, low levels of a protein normally expressed in RBCs, PNP, were linked with the presence of bone metastasis.

The correlation of RBCs protein levels with clinical variables led to the identification of the Lysosome-associated membrane glycoprotein 2 (LAMP2) to be overexpressed specifically in RBCs and higher levels associated with shorter outcomes in metastatic BC patients. Accordingly, with our data, LAMP2 expression levels could act as a reliable biomarker for diagnosing of metastasis in BC, and its specificity and sensibility are increased when it is combined with the blood test parameters which account for RBCs status, haematocrit and, RDW. Interestingly, the capability of these parameters to diagnose metastasis in this patient cohort themselves is lower than LAMP2 alone (data not shown).

LAMP2 is an important regulator in the effective maturation of both autophagosomes and phagosomes and has been involved in cell survival in BC(45). In line with this, the most represented cellular composition of the differential patients’ proteins identified in this work are lysosomes, mitochondria, and extracellular vesicles (EVs). Under normal physiological conditions, RBC-derived EVs constitute 7.3% of EVs in whole blood, indicating that RBCs are one of the main sources of EVs in peripheral blood. Several studies have shown that EVs play key roles in cell-to-cell communication and in cancer they can regulate metastasis tropism. In addition, EVs are the main miRNA carriers in the circulatory system. miRNAs play regulatory roles in the terminal differentiation process of RBCs and they accumulate in mature RBCs(46). Thus, the presence of a higher quantity of EV-related proteins in RBCs from patients could be a result of an increase in EVs accumulation due to the hijack of the EV signalling network during tumor progression.

One limitation of this study is that the GO analysis should be read cautiously since it includes the set of proteins newly identified in RBCs that are yet to be included in the databases as RBCs related. Human RBCs proteome has been previously described by different groups using diverse technologies(9,10,47). However, the LC-MS/MS in DDA mode performed in the present study, combined with a SWATH-MS quantitative, can give a complete proteomic profile of RBCs and also give quantitative values of the proteins. Therefore, the SWATH-MS technology used in this study is a “label-free” quantitative proteomic technique more sensitive and with higher accuracy than those used previously, giving more precise results. Importantly, this work has identified 50 novel proteins in HD RBCs, thus it is necessary to include them on the RBCs databases to be up-to-dating in hand with the new technology developments.

During the metastatic cascade, there are micro-environmental and systemic processes that contribute to cancer regulation, such as immune surveillance(48). RBCs are now considered cells with systemic influence and a dynamic relationship with their environment(11). RBCs circulate in plasma together with a diversity of cells, therefore, many alterations within and on RBCs may result from contact with plasma proteins or soluble factors (including drugs), with substances released from activated cells and, likewise with nucleated cells as CTCs and other circulating tumor material. Although RBCs were historically considered inert to regulatory signals from other cells, they are well equipped with the machinery required for intercellular communication. Thus, as recently published, are capable to regulate the biological processes of neighbouring cells, becoming a novel regulatory cell(12). The observed changes in RBCs from BC patients compared with HD are probably multifactorial where the tumor microenvironment may be having a part, together with inflammation. However, more research is needed in this direction. Remarkably, RBCs are easy to access, highly abundant, and a systemically distributed biological component(11), thus, RBC proteins can be useful biomarkers for cancer monitoring of BC patients and potentially of other tumor types and may constitute a new kid on the block in the liquid biopsy field.

## Methods

### Patients and samples

Blood samples and associated clinical information were obtained at the University Hospital Complex of Santiago de Compostela (Spain). All patients and HD gave written informed consent. All samples were anonymized. The study was conducted according to the guidelines of the Declaration of Helsinki, and approved by the Ethics Committee of Galicia approval reference number 2015/772.

In total, 126 blood specimens from 80 BC patients and 46 non-cancer donors (healthy donors, HD) were collected for this study with an average age of 57 years (29-83years). The median age among patients and donors was similar (t-test, p > 0.05). The HD had different illness including diabetes (n=4), hypertension (n=5), fibromyalgia (n=4) and/or osteoarthritis (n=3). Patients’ clinical information is summarized in Table S1.

### Samples preparation

10 mL of blood (EDTA tube) were centrifuged (1700 g/10’) to isolate RBCs after discarding plasma and Peripheral Blood Mononuclear Cells (PBMCs). RBCs were lysed in 40 mM HEPES (Sigma-Aldrich, St. Louis, MO, USA), 2 % Triton-x100 (Sigma-Aldrich, St. Louis, MO, USA), 200 Mm NaCl, 40 mM MgCl_2_, 20 mM EGTA (Sigma-Aldrich, St. Louis, MO, USA), 80 mM β-glycerophosphate (Sigma-Aldrich, St. Louis, MO, USA), and centrifuged to obtain supernatant containing total protein. Total protein quantitation was performed with DC Protein Assay following the manufacturer’s instructions (Biorad, CA, USA).

One additional EDTA tube was used for blood test analysis and processed in the clinical laboratory of the Oncology Department by standard assays.

### Proteomic analysis by TripleTOF 6600 LC-MS/MS system

#### Protein digestion

To make global protein identification, an equal amount of protein from RBCs of 86 samples (BC patients n=53, HD: n=33) was loaded on a 10% SDS-PAGE gel to concentrate the proteins in a band. The band was processed as described previously(49).

*Liquid chromatography-tandem **mass spectrometry** (LC-MS/MS) in data-dependent acquisition (**DDA**) mode-shotgun analysis.* Digested peptides of all individual samples from RBCs were separated using Reverse Phase Chromatography as described previously(49).

*Protein quantification by SWATH-MS analysis* (Sequential Window Acquisition of all Theoretical Mass Spectra). To build the MS/MS spectral libraries, the peptide solutions were analyzed by a shotgun data-dependent acquisition (DDA) approach using micro-LC-MS/MS as described(49). The MS/MS spectra of the identified peptides were then used to generate the spectral library for SWATH peak extraction using the add-in for PeakView Software (version 2.2, Sciex, MA, USA), MS/MS ALL with SWATH Acquisition MicroApp (version 2.0, Sciex, MA, USA). Peptides with a confidence score above 99% (as obtained from Protein Pilot database search) were included in the spectral library. For relative quantification by SWATH-MS analysis, SWATH-MS acquisition was performed on a TripleTOF 6600 LC-MS/MS system (Sciex, MA, USA). Peptides from RBCs samples from all HD and BC patients were analyzed using the data-independent acquisition (IDA) method making 3 technical replicates per sample as described previously(49).

### ELISA assay

Total RBC protein, extracted as previously mentioned, was used for the determination of LAMP2. The ELISA assay (Company ABclonal, Inc., MA, USA) was performed following the manufacturer’s recommendations using a 1:100 dilution. LAMP2 concentration values were normalized with paired haematocrit values.

### Statistical analysis

Statistical analysis was performed using GraphPad Prism 6.01 software (GraphPad Software Inc.) and R Studio (Version R-3.6.3). Wilcoxon signed-rank test was used for media comparisons and Fisher exact test or Chi-square test for association analysis. Progression-free survival (PFS) and overall survival (OS) were visualized using Kaplan-Meier plots and tested by the log-rank test. Only p values < 0.05 were considered statistically significant.

The normalization and differential expression analysis of the proteomic data were performed using the Bioconductor NormalyzerDE package. The Variance stabilization normalization (VSN) method was chosen among the different normalization options provided by the package. To fit the logistic regression models was used the glm function of the stats package of R. The subsequent analysis of the ROC curve was performed using the pROC package of R.

For Gene Set Enrichment Analysis (GSEA), Gene Ontology (GO) analysis, and protein interaction the used tool was GO-Shiny V0.741 or String (https://string-db.org/), considering strength > 1.2, > 3 proteins and false discovery rate (FDR) < 0.05. Venn diagrams were performed using http://www.interactivenn.net/(50).

## Data Availability

The datasets generated and/or analyzed during the current study are available in PRIDE. The submission of the partial ProteomeXchange was done in accordance with the MAIPE standards.

## Supplementary Materials

Table S1. Clinic-pathologic characteristics of the BC patients and healthy donors; Table S2. Summary of numbers of peptides identified from RBCs extracts from pools of healthy donors (HD), M0, and M1 BC patients by shotgun; Table S3-S5. List of peptides of HD, M0, or M1 pooled samples from shotgun analysis; Table S6. Library of proteins identified by SWATH-MS in HD samples; Table S7: List of differentially expressed proteins by SWATH-MS; Table S8: List of novel proteins identified by Swath-Ms or shotgun compared with reported RBCs databases.

## Acknowledgements

The authors would like to thank the patients and healthy donors who had participated in this study. To all the personnel of the Oncology Service at the University Clinical Hospital of Santiago de Compostela, for their help with patient care and sample management, especially to Carolina García and Cristina Blanco. Also to Gloria García, Laura Muinelo, and Miguel Abal for their scientific advice and discussion. This research was supported by AECC (IDEAS18108COST) and Roche-Chus Joint Unit (IN853B 2018/03), funded by Axencia Galega de Innovación (GAIN), Consellería de Economía, Emprego e Industria. CYG is supported by Axudas Predoutorais da Xunta de Galicia.

## Authors’ contributions

The methodology was carried out by TPV, SB, AGT, CYG, CA, MCV, VV, AV, JFC, and PP. TPV, SB, AGT, and CC performed the formal analysis of the results. The conceptualization and supervision were carried out by CC and RLL. TPV and CC wrote the original draft. TPV, SB, ABD, RCP, and CC were involved in the discussion, writing, review, and editing. RLL and CC were involved in the acquisition of funding. All authors have read and agreed to the published version of the manuscript.

## Conflict of interest

R.L.-L. reports grants and personal fees from Roche, Merck, AstraZeneca, Bayer, Pharmamar, Leo, and personal fees and non-financial support from Bristol-Myers Squibb and Novartis, outside of the submitted work. The other authors declare no conflict of interest. A related patent entitled “In vitro method for the diagnosis or prognosis of breast cancer” (PCT/ EP21382448.5) has been deposited.

## The paper explained

### Problem

Currently, no biomarkers with clinical utility for the follow-up and diagnosis of metastatic disease in patients with breast cancer are available. Besides, it is unknown whether circulating tumor cells modify red blood cells during tumor spread.

### Results

In this study, we have characterized for the first time the RBC proteome of patients with breast cancer in different stages of the disease versus healthy donors. We have identified 50 proteins not described so far in RBCs. We also identified a differential proteomic profile in breast cancer patients. LAMP2 come up as a marker with prognostic and diagnostic potential in the metastatic setting.

### Impact

This study revealed that the presence of a tumor modifies the RBC proteome and the value of RBCs proteins in the prognosis and diagnosis of metastatic breast cancer in a non-invasive way. Besides, it has been described novel RBC’s proteins. Our data provide new information that could open a new path of study in the context of disseminated disease.

## Abbreviations

CTCs: circulating tumor cells
DTCs: disseminated tumor cells
RBCs: red blood cells
BC: breast cancer
HD: healthy donor
HB: haemoglobin
EE: extramedullary erythropoiesis
MS: mass spectrometry
PFS: progression-free survival
OS: overall survival
M0: non-metastatic
M1: metastatic

